# Precision of reaches and proprioception in motor control and adaptation

**DOI:** 10.1101/2025.06.14.659693

**Authors:** Denise Y.P. Henriques, Raphael Q. Gastrock, Bernard Marius ’t Hart

## Abstract

How do precision of movement and proprioception influence motor control and adaptation? Several theories—such as the exploration-exploitation hypothesis—propose that variability plays a key role in motor performance and learning. However, empirical measures of motor and proprioceptive precision are often limited by small sample sizes, and proprioceptive estimates, especially those relying on efferent signals, are difficult to isolate and quantify. In this study, we leveraged a large dataset of 270 participants—including a subsample of older adults (ages 54– 84)—to assess the precision of hand movements and proprioceptive estimates, and to examine whether these factors predict individual differences in motor learning and adaptation. We found that baseline reach variance did not predict learning or changes in hand localization. Although active hand localization (which includes efferent contributions) was slightly more precise—showing an 8.6% reduction in variance—this suggests that unseen hand estimates rely primarily on proprioception. Neither motor nor sensory precision varied with age. However, reach aftereffects were modestly associated with proprioceptive precision before training and proprioceptive recalibration after training. No other measure of learning or variance was reliably associated. These findings suggest that reach aftereffects may partly reflect changes in hand proprioception, but overall, we identified no predictors of adaptation to a rotated visual cursor.

## Introduction

What impact does the precision of movements and the noise in state estimates of the arm have on how sensorimotor systems operate and learn? While there have been several studies investigating the role of motor variability on these processes, very few studies have rigorously measured the variability by which we can localize our unseen hand based on proprioception and predictive signals. Here, we analyzed a large dataset of 270 participants, all tested in the same lab using the same robotic setup, who estimated the location of their unseen hand following both self-generated movements (with predicted sensory consequences) and robot-generated movements (without predicted sensory consequences). In other trials, these participants made goal-directed hand movements with and without cursor. This way we can explore the contribution of proprioceptive signals and efferent signals on localization and motor performance. We then tested whether variance in reach and hand localization measures predicted individual differences in motor performance during the rotated session, including the rate and extent of visuomotor adaptation and the size of implicit aftereffects.

### Estimating hand position

Accurately estimating the position of the arm before movement is essential for effective motor planning and execution. When the hand is visible, visual feedback supports this estimation; however, in its absence, the central nervous system relies on both proprioceptive afferents and efferent-based signals. Proprioceptive estimates arise from sensory inputs transmitted by receptors in muscles, joints, and tendons (for a review, see [1]). In parallel, a copy of the motor command—known as an efference copy—is generated during movement planning and used to predict the sensory consequences of the upcoming action [2–4]. These predictive signals are integrated with sensory feedback, enabling a more accurate and robust multimodal estimate of the limb’s state and improving the precision of future motor commands.

Both proprioceptive and efferent-based information have noise that would impact the fidelity or precision of estimates of hand position. Yet, very few studies have rigorously investigated how precisely we can localize our unseen hand, and how in turn this impacts the precision of subsequent movements (e.g.[5–7]). Moreover, how efferent signals contribute to these estimates and motor performance is particularly difficult to measure since efferent-based estimates are accompanied by proprioception. In our lab, we have devised a set of tasks that gauges how well people can estimate their unseen hand based on only proprioceptive information, or based on both proprioceptive and efferent information. In this way, we have attempted to isolate the efferent-based or predictive component by measuring the differences in these two estimates ( [4,8,9]). From this, we attempt to answer a few novel yet important questions. Does the precision of movements and proprioception affect motor control and adaptation? How do sensory and efferent-based sources contribute to estimates of hand position. How does precision in these estimates relate to motor variability, and how do these different sources of noise impact adaptation? And does this change with age?

How proprioceptive and efferent-signals are combined for localizing the hand remains poorly understood, in part because these signals are difficult to measure and model. This study builds on our earlier work ([4,8,9]), using a substantially larger dataset to examine how these signals are integrated. One possibility is that the brain combines these sensory and efferent inputs through a simple weighted average, without considering their relative precision; however, such models fail to account for the observed variability in two previous hand localization studies. A more efficient strategy would be to integrate inputs using a maximum likelihood estimate (MLE), where the brain combines available information based on its reliability to produce the most precise estimate of limb position ([10,11]). While this approach has been supported in studies combining visual and proprioceptive signals (e.g.,[12–17]), it does not always hold ([18–23]). Importantly, there are few studies examining how the brain combines proprioceptive and predictive (efferent-based) signals. Our previous studies [[4,9]] did not support MLE-like integration in this context. In the current study, we revisit this question with a larger sample and increased trial counts per participant to test the robustness of signal integration mechanisms in proprioceptive and efferent-based hand localization.

### The effect of age

Given that our dataset included older adults, we also investigate whether the precision of hand localization estimates varies with age. Some studies suggest that proprioceptive acuity is poorer in older adults than younger adults (see [24] for a review), including one that tests felt hand position in our own lab [25,26]. Yet, the current study includes a much larger sample of older participants (N = 38) and a larger number of proprioceptive trials to gauge this precision.

Moreover, the previous studies, including the one in our lab, measure estimates of unseen hand localization using a judgment or a matching task which can take orders of magnitude longer, leading to substantial decay of any effect we’d like to measure and be less intuitive than merely reaching to the unseen target hand with the opposite visible hand. For instance, we noticed that our older participants have far less trouble doing the localization task for this study than our previous 2AFC task [25–27]. In any case, there are also many studies that do not find an effect of age on proprioceptive acuity ([5–7,28–31]). In the current study, we are also able to measure and compare proprioceptive and efference-based estimates of hand position for older adults and compare them with a large group of younger adults for greater statistical power.

### How does variability in movements and hand estimates impact learning

Several studies propose that higher variance in movement directions increases the speed of motor adaptation (e.g., exploration-exploitation hypothesis, also see [32–34]). Some studies with larger cohorts have found no relationship between motor variability and the speed of motor adaptation [35,36]. While these findings may cast doubt on the notion that higher baseline motor variability reliably predicts faster adaptation, this idea continues to inform several models of motor learning. It’s also worth noting that other sources of variability, such as sensory or state estimate noise, have not been fully explored and may contribute differently to adaptation. Therefore, we put it to the test here with over 100 participants.

Additionally, the speed or rate of learning is often inadequately measured in many reach adaptation studies. Usually this is based on the average reach performance across the first few rotated trials which makes for a maximally noisy estimate of the rate of adaptation. Here we get a descriptor of the rate of adaptation by fitting a standard exponential learning curve to the full adaptation phase. Since that is based on many more trials, any accidental fluctuations due to measurement or execution noise get canceled out and the estimates of the rate of adaptation are minimally noisy.

We test if three measures of adaptation can be predicted: the rate of adaptation, the extent (asymptotic value) of adaptation and the size of reach aftereffects. We look at four potential predictors that are estimates of baseline variance: variance in reaches with regular feedback and in open-loop no-cursor reaches, as well as variance in two kinds of hand location estimates. Since some of the previous work from our lab [9,37,38] has suggested that the magnitude of hand localization shifts (or proprioceptive recalibration) is related to the size of reach aftereffects, we also test, if the predictions improve when variance in hand localization is combined with hand localization shifts to predict reach aftereffects.

## Methods

### Participants

Two-hundred and seventy participants (232 younger adults (152 female) and 38 older adults (19 female)) were recruited from either York University’s Undergraduate Research Participant Pool (URPP), Kinesiology Undergraduate Research Experience (KURE), the Centre for Aging Research and Education (YU-CARE) participant pool, and through acquaintances from the York University community. All participants were right-handed, had normal or corrected-to-normal vision, and were naïve to the purpose of the experiment. Table 1 shows the primary experiment source of all participants who gave informed consent.

**Table 1.**
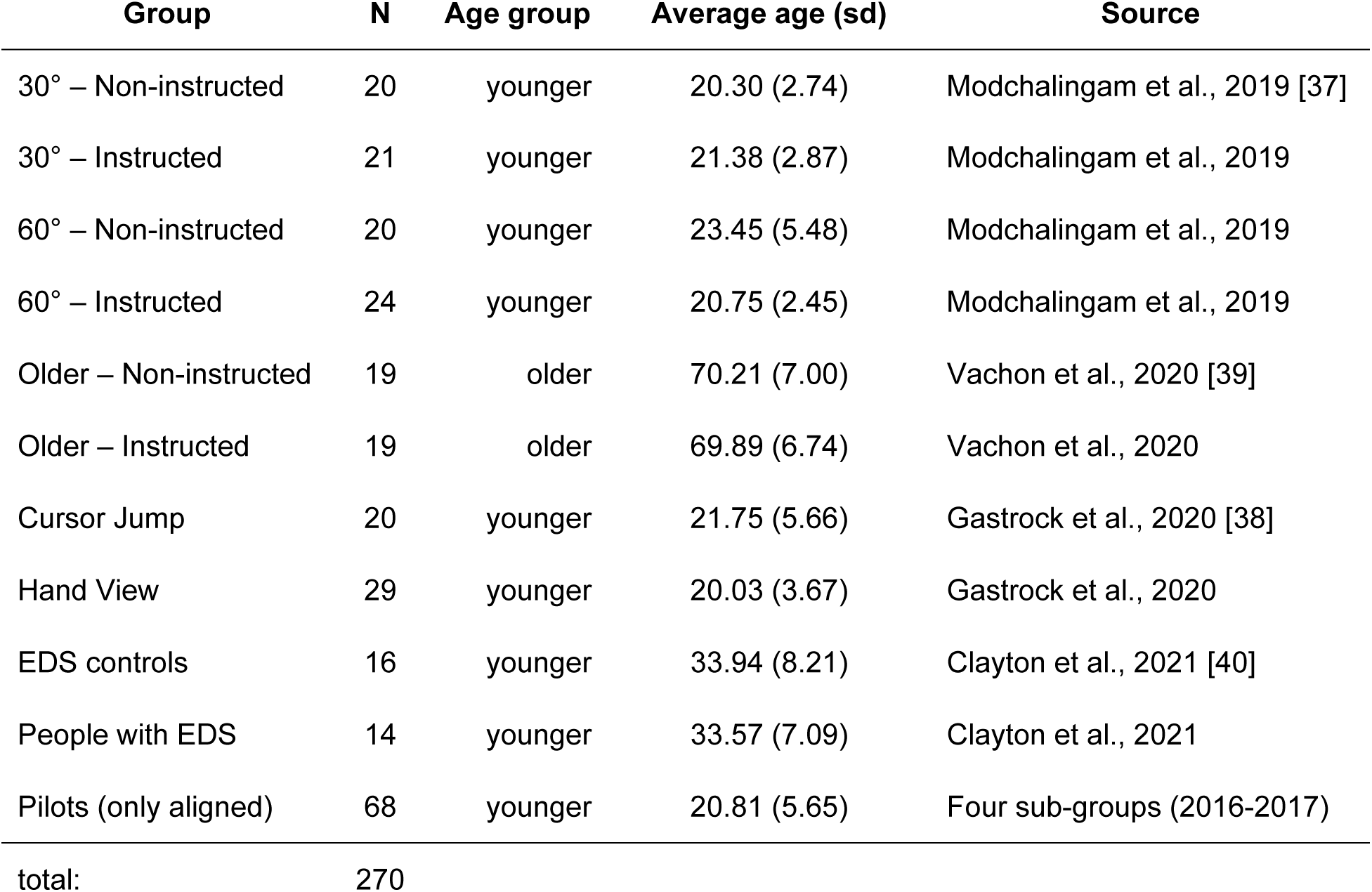
Primary Experiment Source of Participants (N = 270). Note that the initial baseline session across all these studies were identical or highly similar. Older participants were 54 years and older (average age = 70, SD = 6.78). All other participants were younger but at least 17 years old. EDS: Ehlers-Danlos syndrome.

| Group | N | Age group | Average age (sd) | Source |
| --- | --- | --- | --- | --- |
| 30° – Non-instructed | 20 | younger | 20.30 (2.74) | Modchalingam et al., 2019 [37] |
| 30° – Instructed | 21 | younger | 21.38 (2.87) | Modchalingam et al., 2019 |
| 60° – Non-instructed | 20 | younger | 23.45 (5.48) | Modchalingam et al., 2019 |
| 60° – Instructed | 24 | younger | 20.75 (2.45) | Modchalingam et al., 2019 |
| Older – Non-instructed | 19 | older | 70.21 (7.00) | Vachon et al., 2020 [39] |
| Older – Instructed | 19 | older | 69.89 (6.74) | Vachon et al., 2020 |
| Cursor Jump | 20 | younger | 21.75 (5.66) | Gastrock et al., 2020 [38] |
| Hand View | 29 | younger | 20.03 (3.67) | Gastrock et al., 2020 |
| EDS controls | 16 | younger | 33.94 (8.21) | Clayton et al., 2021 [40] |
| People with EDS | 14 | younger | 33.57 (7.09) | Clayton et al., 2021 |
| Pilots (only aligned) | 68 | younger | 20.81 (5.65) | Four sub-groups (2016-2017) |
| total: | 270 |  |  |  |

### Consent to participate

The experimental protocol was approved by the Human Participants Review Committee at York University, in accordance with the ethical standards set forth in the Canadian Tri-Council Policy Statement: Ethical Conduct for Research Involving Humans (TCPS-2) and the Declaration of Helsinki. All participants provided written informed consent prior to participation in the study.

This study was funded by a NSERC Discovery grant to DYPH.

### Apparatus

Participants sat on an adjustable chair facing a semi-reflective screen located 14 cm above a two-joint robot manipulandum, which reflected images displayed on a computer screen allowing them to be projected on the same horizontal plane as the manipulandum (Figure 1A). Using their right hands, participants held a vertical handle of the robot manipulandum with their thumbs resting on top of the modified handle (Interactive Motion Technologies Inc., Cambridge, MA, USA). Visual stimuli were presented from a monitor (Samsung 510 N, refresh rate 60 Hz) located 28 cm above the robotic arm. A touchscreen (Keytec Inc., Garland, TX, USA) was horizontally mounted 2 cm above the robotic arm to record reach endpoints made towards proprioceptive targets.

**Figure 1.**
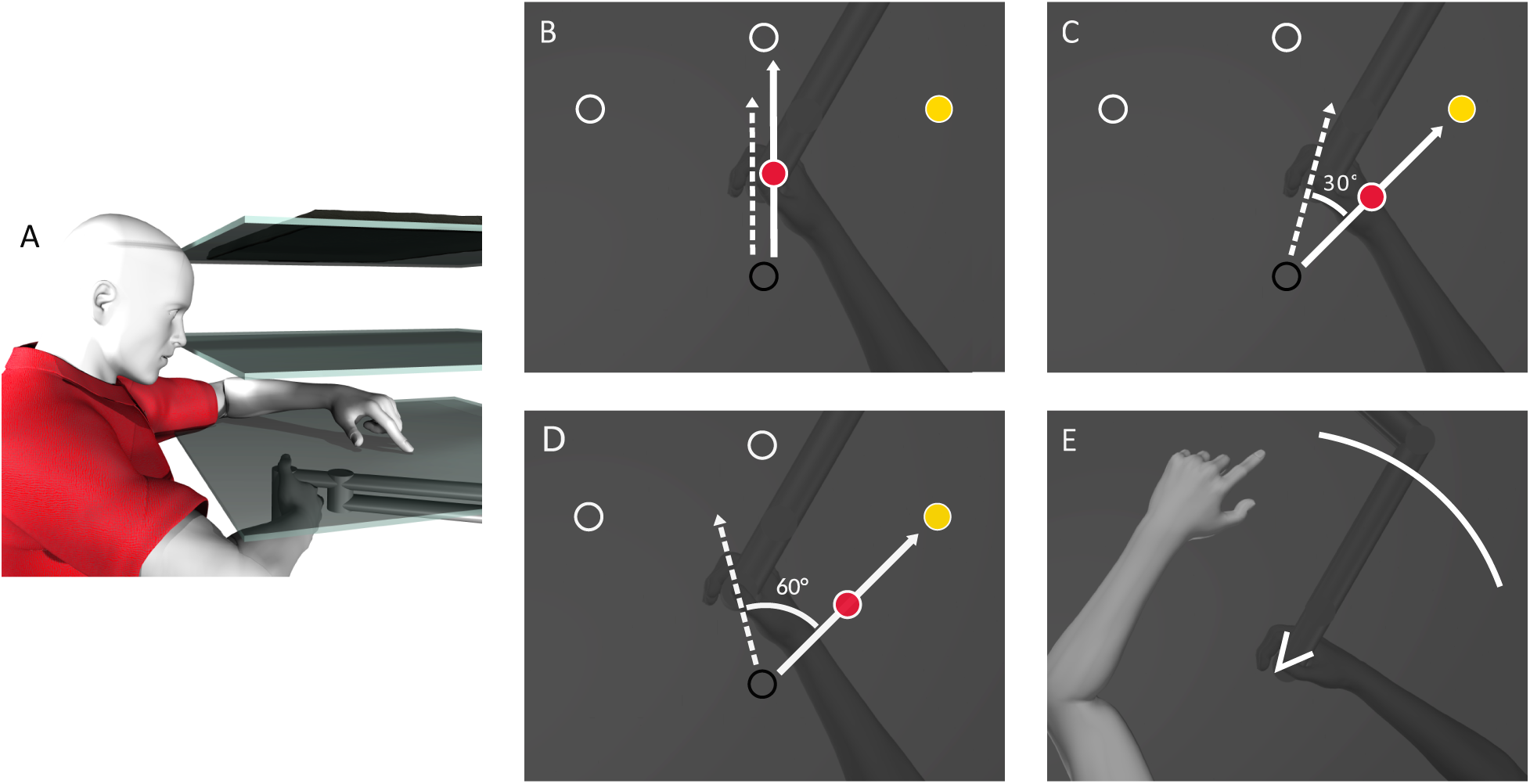
Experimental setup and procedure for most of our experiments adapted from [37]. **A.** Participants held a robot manipulandum with their right hand beneath a touchscreen (bottom surface), while viewing a reflective screen (middle surface). Visual stimuli were projected from a monitor positioned above the reflective screen (top surface) and appeared aligned with the hand workspace. During localization trials, participants used their visible left hand to indicate the position of their unseen right hand on the touchscreen panel. **B-D:** On a given trial, participants reach to one of three targets (white circles/yellow disc) using a hand-cursor (red disc) as quickly and as straight as possible. The cursor is aligned with the unseen hand (**B**) for all studies, or rotated 30° (**C**) or 60° (**D**) depending on the study and group. **E.** During localization tasks, participants used their visible left hand to indicate where they crossed an arc with their unseen right hand. For participants in the 60° groups, a V-shaped wedge of 60° served as an indicator as to which part of the workspace to move their unseen hand. See original manuscripts for more specific details.

Participants completed the task in a room with very low ambient light. The view of their trained right hand was completely occluded by the reflective surface (Figure 1A) for all but the training trials in one group [38].

### Experimental tasks

Data comes from 14 experimental groups (Table 1) that share near-identical hand localization tasks (active and passive hand localization), which were interspersed within reaching trials to a visual target using a cursor that was aligned with the hidden, right hand (aligned training). This aligned session is immediately followed by a rotated session with a rotation of 30° or 60° under various conditions. We begin our analysis using data from the aligned session, which includes proprioceptive estimates and simple, unperturbed reach movements. Although this session is typically treated as a baseline, here we use it to gain new insights into how proprioceptive and efferent signals contribute to hand localization in the absence of perturbations. To our knowledge, such alignment-based analyses have not previously been used to probe the integration of perception and action. We then extend our investigation to examine how the precision of these hand localization estimates relates to subsequent motor adaptation—addressing novel questions about whether variability in perception predicts the rate or extent of learning. Figure 2 shows the shared session structure across all 14 independent experimental groups. The datasets generated and analysed during the current study are available in the following repository: https://osf.io/dhk3u/.

**Figure 2.**
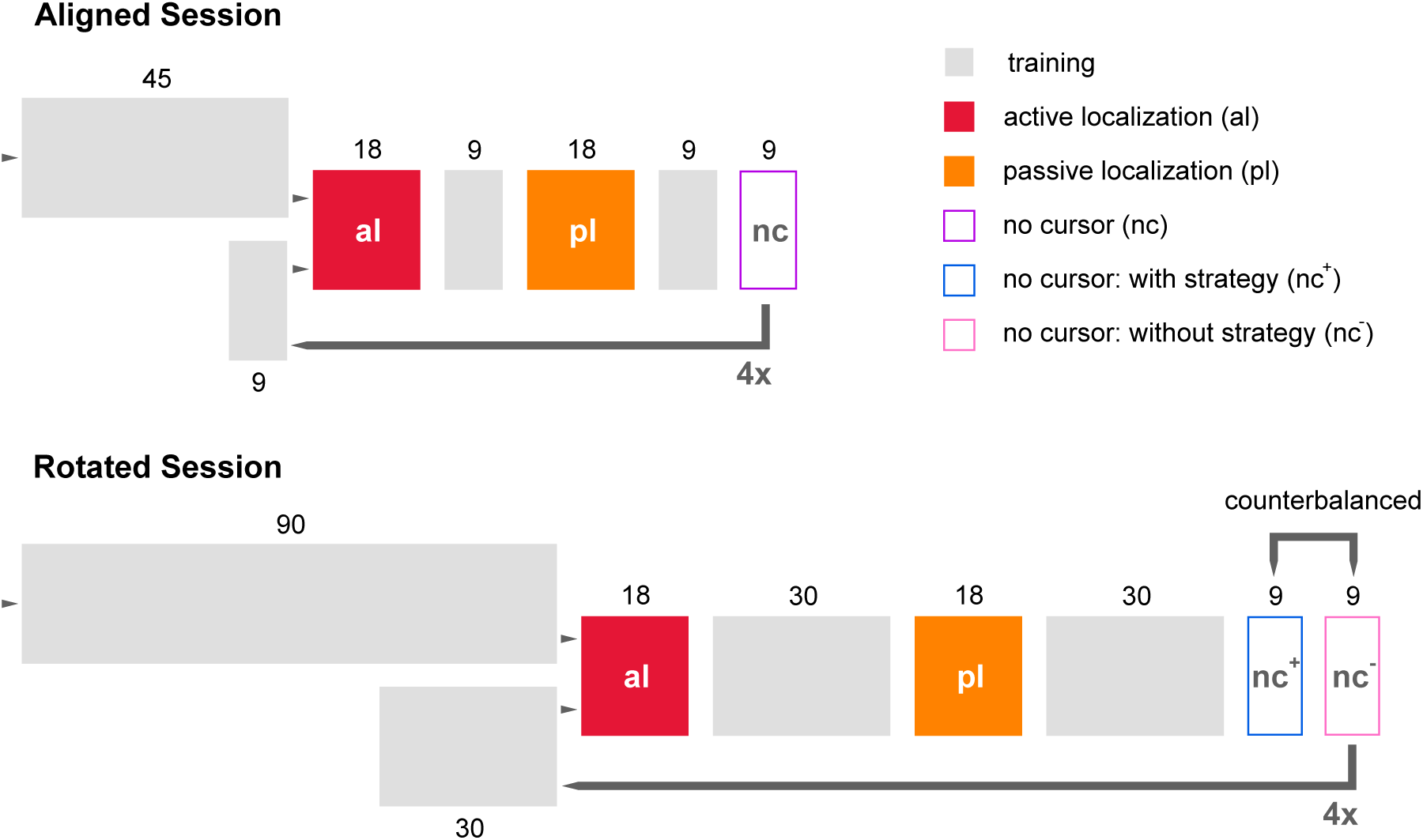
Session sequence. All experiments used in our analysis followed the same schedule starting with a baseline (aligned) session (top), where blocks of active and passive localization trials occurred between reaching trials to visible targets with a cursor that was aligned with the hand. This was followed by a rotated session (bottom) that had longer blocks of reaches with the cursor, and two types of no-cursor trials (we use only one here). Grey blocks represent reaching trials with visual feedback of the hand cursor. Active localization trials are depicted in red blocks, while passive localization trials are depicted in orange. No-cursor trials, excluding strategy, are depicted in hollow purple blocks. The blue block indicates no-cursor trials in which participants were asked to include their strategy; these were not used in the present analyses. This identical first baseline session was immediately followed by the rotated session (bottom panel) where the cursor was rotated by either 30° or 60° clockwise (see Table 1). The schedule for the rotated session was consistent across groups, except that some participants received explicit instruction about the cursor rotation beforehand. For details on a variation in how the visual perturbation was presented, see [38].

#### Localization tasks

To address our main questions, we measure the accuracy and variability of estimates of the location of the robot-displaced or self-generated hand at the end of the movement, in the baseline session and after visuomotor adaptation. During the two localization tasks, the unseen right hand served as the target, where key differences lie in how the target-hand achieves its final position (Fig 1E). In the active localization condition (Fig 2, red box), participants held the robotic manipulandum with their unseen hand and made volitional movements in a direction of their choosing within the boundary of the arc or v-indicator as illustrated in Fig 1E. When participants moved their unseen right hand 12 cm from the home position, a force was applied to gently stop the hand and then lock it in place, preventing further movement and maintaining that hand location for 700 msec. They then returned to the home position along a constrained path at their own pace. Using their visible, untrained left hand, participants indicated on a touchscreen where they perceived their unseen right hand to have been at the point of contact. The constrained-return path used in this and all trials in order for the unseen hand to quickly return to start position was generated by a perpendicular resistance force of 2 N/(mm/s) and a viscous damping of 5 N/(mm/s).

In the passive localization condition (Fig 2, orange box), the robot manipulandum moved the participant’s hand to a selected location 12 cm away from the home position .These selected locations matched those that participants had self-generated during the preceding active localization condition, where they voluntarily chose where to place their hand. After the passive movement, participants returned their hand to the home position along the constrained-return path. Using their visible, untrained left hand, participants indicated on the touchscreen the perceived location their unseen right hand had been passively moved to during the outward reach. Localization data were collected both during training with veridical feedback, in the aligned / baseline session (Fig 2, top row) and following training with rotated feedback (bottom row). We collected 72 hand localization trials in both active and passive localization in the aligned session, and an equal amount in the rotated session, for 288 hand localization trials in total per participant.

In both active and passive localization tasks, participants first moved their hand away from the endpoint of the reach, back to the home position, such that the current proprioceptive signal could not be able to fully overrule any efferent signals. They were then instructed to indicate where their outward movement had crossed the white arc, that is, the instructions focussed on the movement, such that efferent signals received some more attention. This organization of the trials was chosen when we found a larger difference between active and passive localization shifts with delayed responses as compared to reporting hand location with the hand still at the endpoint of the reach [4,9,41]. With this method, efferent signals are sure to contribute to the final estimate, which is necessary to estimate its relative contribution to hand location sense.

To ensure that hand localization reflected only non-visual estimates, the trained right hand was kept completely out of view beneath the touchscreen, with a black cloth draped over the participant’s shoulders to block visual feedback. During localization trials, participants used their visible, untrained left hand—illuminated by a small lamp—to indicate the perceived location of their unseen right hand after returning to the home position (Fig. 1E). No visual targets were ever presented during these trials, as doing so would have biased hand position estimates. This design prevented confounds from visual guidance, interlimb transfer, or feedback from the right hand at the end of the reach, ensuring that localization reflected proprioceptive and/or efferent-based estimates alone.

#### Training with aligned feedback (baseline)

To assess the relationship between motor precision and hand-estimate precision, we also measured reach variance across our series of experiments. During aligned training (Figure 1B, Figure 2, gray boxes), participants received visual feedback of their hand position via a continuously displayed green cursor (1 cm in diameter). In one group only, participants saw their hand along with the aligned cursor ([38]) during the training trials. After participants placed their hand at the home position for 300 ms, the target appeared. Participants were asked to reach targets as accurately and as quickly as possible. Trials where participants received visual feedback of the cursor would end when the centre of the cursor was within 0.5 cm of the target’s centre. After reaching the target the unseen right hand was held still for 500 ms. At this point, the target disappeared, which signaled to the participant to return their hand to the home position along a constrained return path. During no-cursor trials (both in baseline and rotated sessions), when participants believed that they had acquired the target, they held their hand in place for 500 ms which indicated the completion of the reach and followed by the disappearance of the target. Participants then moved the robot handle back to the starting position along a constrained path, similar to trials with visual feedback of the cursor.

#### Training with rotated feedback

To address our second set of questions, we included those participants in the dataset who also adapted to a visuomotor rotation (7 of the experimental groups) following baseline conditions. This rotated session was similar to the aligned session, except the cursor motion was misaligned from the hand by either 30° clockwise (CW) rotation (Fig 1C), or a 60° CW (Fig 1D); and we introduced more trials with the rotated cursor to allow for complete and sustained adaptation (Fig 2, bottom row).

#### Clarifications on Task Demands, Kinematics, and Measurement Choices

Because the present study compares variability across perceptual and motor tasks, we provide additional detail on several task features that may influence movement kinematics or hand-localization estimates. First, although perceptual and motor measures necessarily differ in certain demands, we ensured that key aspects of their timing and general movement requirements were comparable, and we avoided task conditions known to artificially increase variability. Participants were free to hold their hand at the final position for as long as they wished in all tasks after the initial 500 and 700 msec hold. For reaching trials, the movement ended once the hand was briefly held steady—after cursor–target overlap for cursor reaches, or after the hand moved at least 80% of the distance for no-cursor reaches. The same brief stability requirement applied to both reach types, after which the virtual return pathway became available. Although this steady-hold criterion is not essential—“slicing” movements typically produce similar learning patterns for the ballistic portion—we used it to ensure that participants consistently completed and stabilized their reaches. Participants were encouraged to “move quickly,” as is typical in visuomotor-adaptation studies, but this instruction was not enforced and carried no penalty; movements were performed at comfortable, self-selected speeds. No speed instructions were given for the left-hand indication movements either, allowing participants to focus on accurately placing their visible left hand at the perceived location of the unseen right hand. Thus, any errors in this task primarily reflect perceptual estimates rather than movement constraints.

With respect to analysis choices, no-cursor reaching movements are generally very straight, making angular deviation at peak velocity and at endpoint highly similar. In our earlier publications using portions of this dataset, we used endpoint measures for comparing aftereffects to localization. In the present work, to maintain consistency between cursor and no-cursor reaches and to enable additional comparisons (e.g., with learning-rate measures), we use angular deviation at peak velocity for both. Importantly, using either measure yields the same qualitative pattern of results.

The number of trials in each experiment was originally determined to address our prior research questions while keeping the overall session duration short enough to avoid fatigue. These studies were not designed specifically to optimize estimates of precision. Nonetheless, the number of observations exceeded the common rule of thumb (≈30 trials) typically considered sufficient for reasonably stable variance estimates. In addition, we conducted simulation-based checks to evaluate whether the number of localization trials—particularly given that estimates were pooled across multiple hand locations—was sufficient to yield reliable measures of variability. These simulations confirmed that the available number of trials provided stable estimates of localization precision.

### Data analyses I: Pre-processing and statistical tests

All data preprocessing and statistical analysis were implemented in R 4.3.2. (R Core Team, 2023) which are available as scripts through this repository: https://osf.io/dhk3u/.

For conciseness, we will not re-analyse visuomotor adaptation, or shifts in hand localization here following adaptation; the reader can review the publications listed in Table 1 to verify that participants were able to adapt to the visuomotor adaptation and produced robust reach aftereffects, and shifts in hand estimates for both passive and active localization following adaptation. Figures in the current study also nicely illustrate the magnitude of the robust effects of learning.

Reach deviations in training trials (aligned or rotated) were taken as the angular difference between the direction of the target and the direction of the hand’s reach at the point of maximum velocity. Reach deviations in no-cursor trials were taken as the angular difference between the direction of the target and the direction of the end point of the hand’s outward reach. Localization estimates were determined as the angular difference between the endpoints of the unseen right hand and each participant’s perceived hand position as indicated on the touchscreen. This reduces each key measure to a single angular dimension.

#### Exclusion criteria

Localization responses were first screened manually for each of the four previous publications (including the pilot data) for obvious response errors (accidental taps on the edge of the screen, etc). We used two additional criteria here. First, the angular deviation of the unseen hand had to be within 45 degrees of the centre of the arc, since participants were instructed to tap on the arc. Second, localization errors had to be within 2 times the interquartile range for each participants’ four subsets of localization responses. Training and no-cursor reaches were also screened manually, and the relevant part of the trajectory was selected.

Given that the cursor was rotated CW, localization shifts should also be in the CW, or negative direction, while asymptotes and reach aftereffects should be in the CCW, or positive direction.

#### Variances in reaches and hand localization

For the training and no-cursor trials, we can calculate a bias for every target - since there usually are systematic biases depending on direction in the work space. We can then subtract those biases so they do not artificially increase our estimate of the standard deviation. So the approach for targeted reaches is to subtract the average bias for each target before determining the overall variance using this de-biased data.

Since localization reaches are made without a target, we can not use that exact approach for localization. But we can estimate the biases across the workspace differently—by fitting a smoothed spline (9 nodes, smoothing factor of 0.5). A spline is a smooth, flexible curve used to approximate data, and in this case, it allows us to model how bias changes as a function of angle. For every participant we base one spline on the aligned active localization trials and one on the aligned passive localization trials. These serve as estimates of a direction dependent mean bias for both the aligned and rotated, active and passive localization (details in [4,9]). After subtracting the bias predicted by the spline on each trial from the localization response, we can determine a de-biased estimate of standard deviation, comparable to those of the no-cursor and training trials.

Then, to obtain baseline variability for direction of hand localization (relative to home position), we calculated the standard deviation of localization responses for every participant, for each localization condition (active and passive). That is, for reach-based angular deviations as well as localization-based angular deviations, we first get an estimate of direction-dependent bias and remove this before determining a measure of variance.

#### Testing differences between variables

We test for differences between measures of variance using regular t-tests in one case, as well as a Bayesian ANOVA and follow-up Bayesian t-tests. Reported Bayes factors reflect the ratio of how likely the alternative hypothesis (there is a difference) is over how likely the null hypothesis (there is equivalence) is, given a non-informative prior and the data. With BF_10_ = 1 both are equally likely. Within the interval ⅓ to 3 (either hypothesis is up to 3 times more likely than the other) there is only anecdotal evidence, and thus no real effect [42,43]. However, a BF_10_>3 or BF_10_<⅓ (or 0.333) is considered moderate evidence in favor of the alternative hypothesis or the null hypothesis, respectively, whereas values of BF_10_ > 10 or BF_10_ < 0.1 are considered strong evidence. Note that in contrast to classic null-hypothesis significance testing (NHST), Bayesian statistics allow confirming a null hypothesis.

#### Testing relationships between variables

For all other analyses, we perform two types of regressions between pairs of variables. For tests with a clear or hypothesized directionality between the two variables (dependency of one variable on the other) we use regular regressions (ordinary least squares regression, OLS). This type of regression minimizes errors between the line of best fit and actual data points along the axis of the predicted variable. However, for situations where we assume no directionality we use orthogonal distance regression (also called PCA-based regression). The implementation here is based on the odregress function in the pracma package by Hans W. Borchers. This type of regression minimizes the errors taken in a direction orthogonal to the line of best fit.

#### Maximum Likelihood Estimate

In order to determine whether efferent and afferent information can be combined in the brain using a Maximum Likelihood Estimate (MLE) in principle, we analyze baseline variability, as a measure of uncertainty, of localization estimates during passive localization and that of active localization. The assumption is that the presence of efferent information would make active localization more reliable than passive localization. We first simply test this with statistics, i.e. a paired sample t-test (or using a Bayesian version of the same test, which -unsurprisingly-gives the same result).

We also did a “reverse” MLE to estimate the isolated reliability of efferent contributions to hand localization on the group data. Reliability of hand localization information when both signals are present is given by:

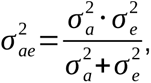

Where 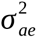 and 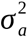 are the measured variance in active and passive localization, respectively. To get the variance in the efferent signals only, we rewrite this as:

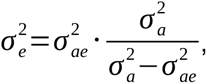

The efferent and afferent signals are then combined as a weighted average, where the weight is the inverse of the relative variance, i.e. the relative reliability of each signal *i* compared to all *n* signals:

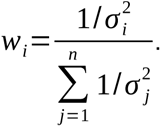

The actual averages of efferent and afferent contributions would be very close to zero, in the aligned conditions, and hence not very informative. However, the relative weights of both efferent and afferent signals are sufficiently informative here.

#### Predicting measures of motor adaptation

We test how well four descriptors of motor adaptation from the rotated session can be predicted with variance measures from the aligned session. These are the speed of learning, the extent of learning, implicit reach aftereffects and shifts in hand localization. The same biases calculated to remove the effects of bias from estimates of variance are also subtracted from measures from the rotated reach deviations and localization responses to get a difference between the aligned and rotated session.

To obtain estimates of the learning rate and extent of motor adaptation, we fit a single exponential learning curve ([41,44]) to reach deviations from the first 90 rotated training trials, predicting the reach deviation *x* on each trial *t*.

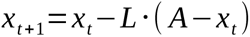

This function has two free parameters: a learning rate L and an asymptote A. The learning rate is restricted to the range [0,1]. and the asymptotic level is restricted to [0,2*max(reach deviation)]. We use a grid search, take the 10 parameter sets that result in the lowest mean squared errors (MSEs) and use those as the starting point for an optimization procedure that minimizes the MSE further. From those we use the parameters from the resulting in the lowest MSE. This is repeated for all suitable participants.

As can be expected, the asymptotes of the exponential fits, which reflect learning extent in degrees, were larger for those participants who adapted to a 60° compared to 30° rotation. So for determining whether motor or sensory variability could predict learning rate, we do not include groups that did 60° rotations. We also don’t include groups that received instructions (inflated learning rates) and can’t include the pilot groups (no rotated phase). This leaves data from 118 participants who adapted to a 30° misaligned cursor when testing whether variance in hand estimates could predict speed of learning.

For testing implicit reach aftereffects, we use the average “exclude strategy” no-cursor reach deviations for each participant (minus the aligned no-cursor reach deviation biases). Since no effect of rotation size, instruction, age or feedback type was observed on these in our previous papers, we use data from all 202 participants who did a rotated phase for these analyses.

We use the magnitude of active as well as passive hand localization shifts (“proprioceptive recalibration”) here as well. Both to test if these are related to the variance in aligned hand localization and if this is related to implicit reach aftereffects.

### Data Analyses II: Addressing research questions

#### The effect of age on localization variability

Given that some of the participants in this study were collected for a study investigating the effect of age on adaptation and changes in hand localization, we took this opportunity to investigate whether proprioceptive variability in both active and passive localization as well as reaches with and without cursor feedback, in this subset of older participants (age: 54-84) differed from that of a large cohort of young adults (17-40). We test this as part of a Bayesian ANOVA with age group as one factor, and the four types of trials as the other factor.

This analysis is followed up with Bayesian t-tests comparing variability in each of the four measures with all other measures. The small difference between variability in active and passive localization is then probed further using reverse MLE described above.

#### The role of localization variability on motor performance and adaptation

One of our main questions was to investigate whether variability of hand location estimates could predict motor variability when reaching with a cursor and without a cursor. Another related question was whether either this location-estimate variability or motor variability could predict speed of adaptation and reach aftereffects.

#### The effects of hand-localization variability on shifts in hand localization

Likewise, to test whether more variable estimates of hand position in aligned training leads to larger changes in hand localization after rotated training, we ran a simple linear regression model on hand localization shift with as the predictor baseline localization variability in both active and passive conditions. To do so, we computed the mean localization shift (i.e. shift in the felt hand position following visuomotor rotation) for each participant for active and passive conditions.

#### The relationship between shifts in hand localization and reach aftereffects

Lastly, we also tested whether visually-induced shifts in estimates of unseen hand location for both active self-generated hand displacement and passive displacement could predict magnitude of reach aftereffects. While this kind of analysis has been done before in the individual studies that make up this database, we ran an orthogonal distance regression in the current paper with the larger database of 270 participants. We also compared the strength of this linear relationship when hand shifts were produced during active localization and passive localization.

## Results

### Movement and proprioceptive precision

While not the main point of our paper, people were quite good at moving their hand radially, represented by an aligned cursor, to a visual target. Accuracy for the estimates of the unseen hand position is illustrated in Fig 3. Younger adults (A-B) do not appear very different from older adults (C-D). As will be described further, estimates of the direction of final location of our unseen hand were also quite precise both when the hand is passively displaced by a robot (Fig 2, red box) or self-directed by the participant (Fig 2, orange box). We can also see direction dependent biases (Fig 3, right side), which we remove as described earlier.

**Figure 3.**
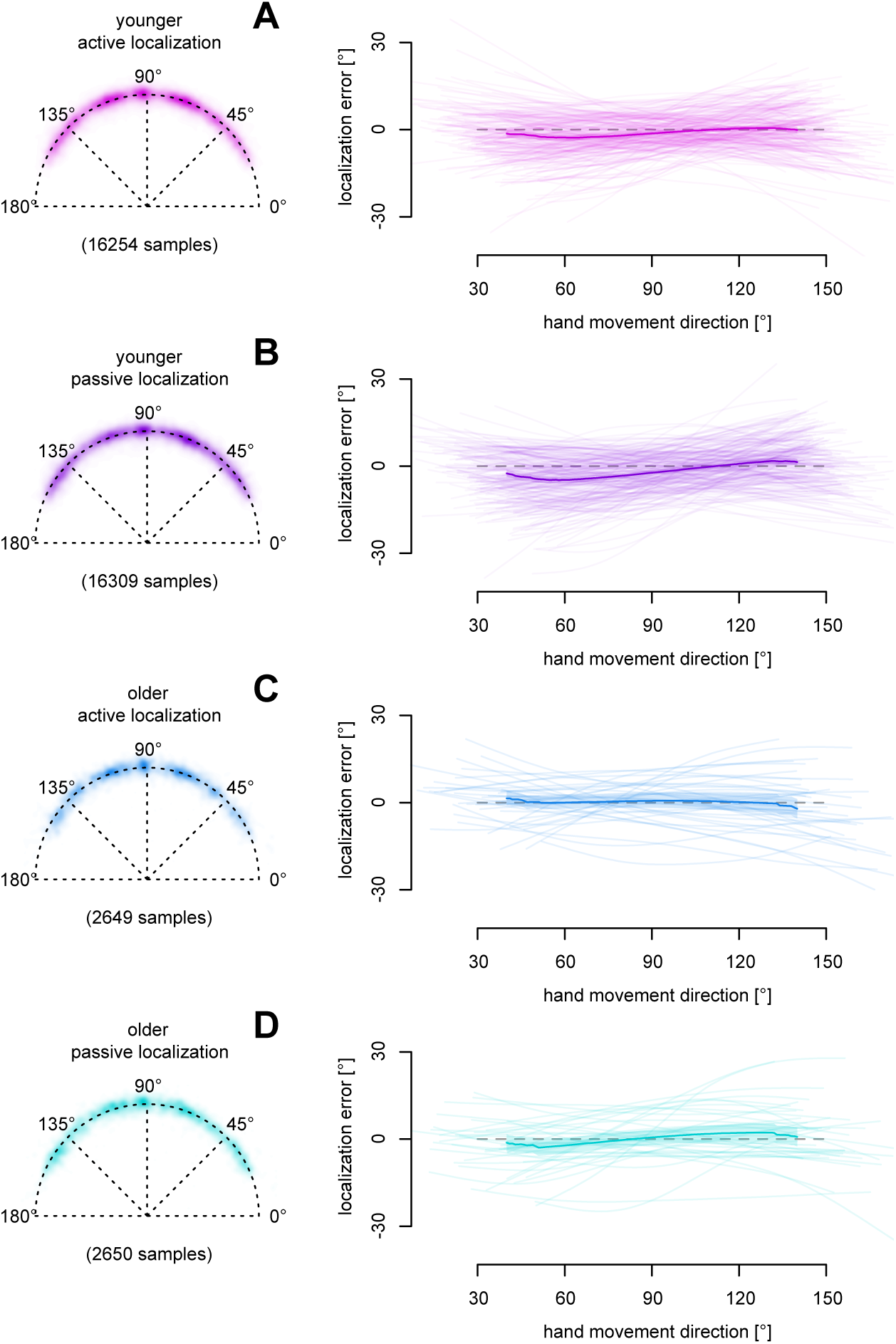
Hand localization performance in the aligned session. Left: an above view of two-dimensional endpoints. Right: “smoothed” estimates of the angular localization error or biases throughout the workspace with light lines for individual participants, and a darker line plus shaded region indicating the group average and a 95% confidence interval over participants. The results for hand-localization after the right hand was actively placed by the participant (**A**, **C**) and passively placed by the robot (**B**, **D**) in young (**A**, **B**) and older (**C**, **D**) adults. Negative angular localization errors are in the clockwise direction. Note that aligned localization errors are on average all fairly low, depend somewhat on the reach direction, and do not appear different between age groups or between active or passive localization.

The magnitude of variance, plotted in Figure 4, differed significantly across the measures of reaches and unseen hand estimates. Specifically, Bayesian analysis across the 270 participants provided strong evidence for differences among these measures (BF_10_ = 6.7 × 10^15^). The strong evidence for differences across motor and perceptual variance measures is primarily driven by the increased variability in no-cursor reaches. These reaches were significantly noisier than cursor-guided reaches (BF_10_ = 2.4 × 10^12^) and both types of hand localization estimates (BF_10_ > 9 × 10^8^). As shown in Fig. 4, no-cursor reaches at peak velocity were approximately 26% more variable than the other conditions. In contrast, we found evidence supporting similarity in variance between cursor reaches and both active and passive hand estimates (BF_10_ = 0.154 and 0.072, respectively). Nonetheless, we observed strong evidence for a small but reliable difference between the two hand localization conditions: participants were slightly more variable when localizing their passively displaced hand compared to when they generated the movement themselves (BF_10_ = 197.16). However, this effect was modest—amounting to just a 0.3° or 5% difference, with a Cohen’s d of 0.248.

**Figure 4.**
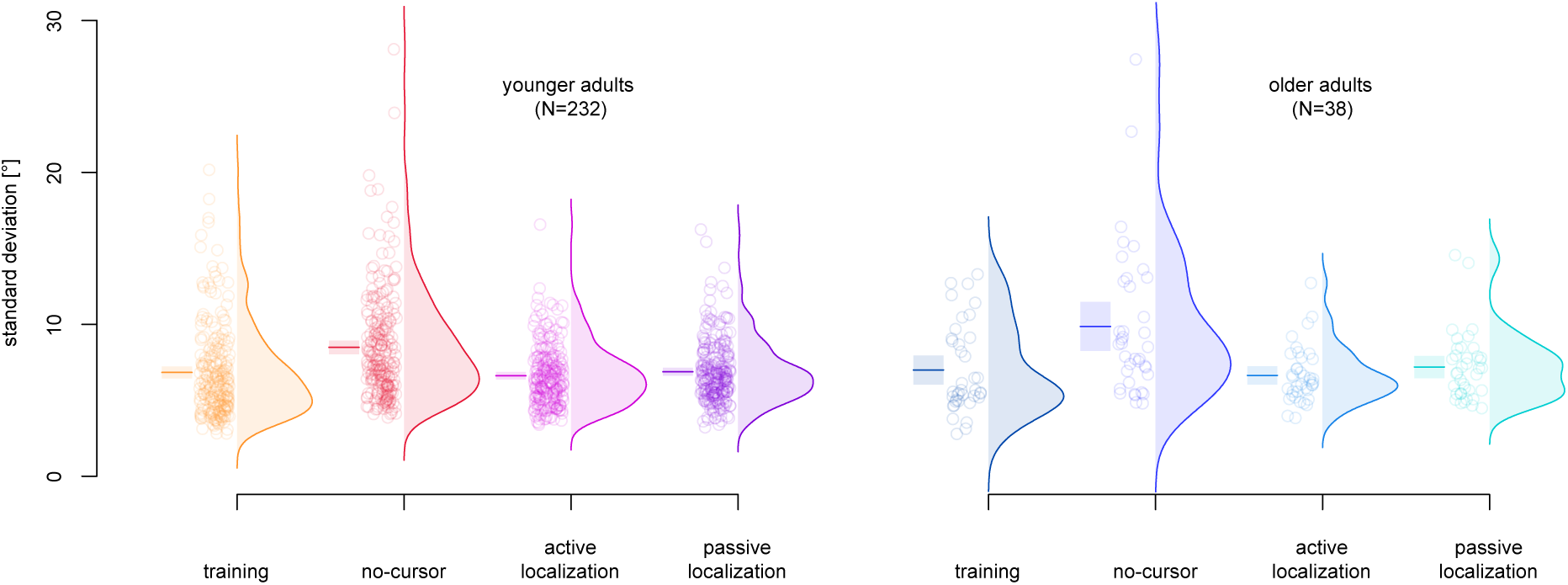
Main measures of angular variation. Standard deviation for reaches made with a veridical cursor (training), open-loop reaches made without cursor feedback (no-cursor), and for estimates of the unseen hand after the hand was actively or passively displaced for younger (left) and older (right) participants. Horizontal lines are the average, displayed within a shaded rectangle denoting the 95% confidence interval. Dots are individual participants. Kernel density estimates are used to visualize the distributions, which all appear uni-modal, although skewed - because of a “floor effect”.

Importantly, this pattern of variance did not differ between age groups. We found moderate evidence supporting the null hypothesis (BF_10_ = 0.13), even after excluding a small subset of participants aged 35 to 40 (BF_10_ = 0.14). Similarly, there was inconclusive evidence for any overall effect of age on these variance measures (BF_10_ = 0.4). Despite our large sample size, there was no indication that older adults were less precise than younger adults in either their reaching movements or their estimates of unseen hand position. In sum, older adults were not more variable than younger adults. Across both age groups, only reaches made without visual feedback showed a marked increase in variability.

Not surprisingly, variability in passive and active localization were highly correlated with each other (shared variance of 67% across all 270 participants) as illustrated in Fig 5. Note this is an orthogonal distance regression since we are agnostic about which serves as the predictee or the predictor in this relationship. This study is one of the first to try to estimate the ‘efferent’ contribution of state estimate of the hand given the difficulty isolating this source of information. Overall, although difficult to quantify, the reduction in variance when both proprioception and efference-signal sources are available compared to when only proprioception is available is not huge, and not consistent across participants with only 59% of participants showing a higher variance for passive hand estimate compared to active ones (dots above the dotted unit slope of Fig 4). Given our large sample size, the difference of 0.3° does get support as a statistically significant effect (t(269)=4.07, p<.001, BF_10_= 197.1). While we cannot directly measure the variance and weight specific to efferent-sources, we can do a reverse MLE calculation (see methods). Since only 59% of participants have a higher variance in passive compared to active localization, we do this across all 270 participants at once, using group average variances. Reverse MLE estimates the weights for afferent signals at 0.914 and 0.086 for efferent signals. This small contribution from efferent signals, either means that state estimates of limb position depend largely on afferent signals such as proprioception, or argues against a straightforward Bayes-optimal integration of efferent and afferent signals in hand localization.

**Figure 5.**
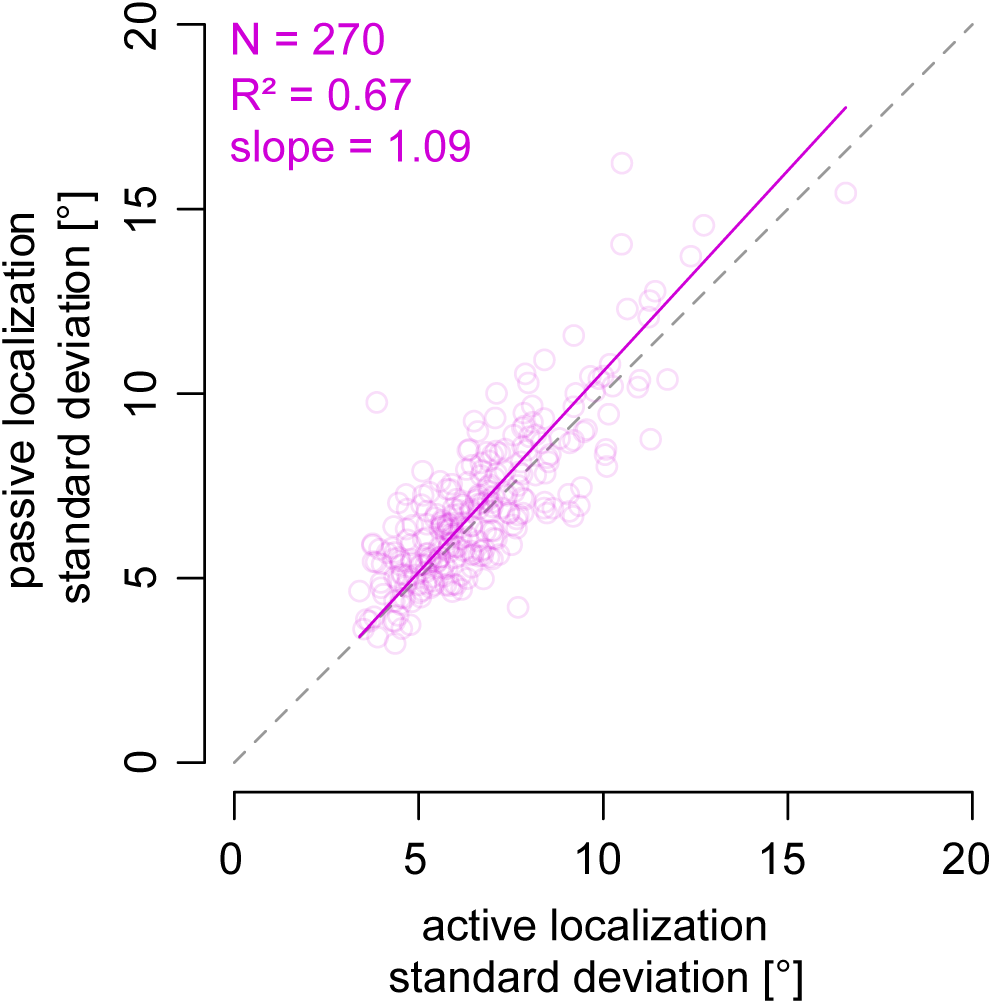
Standard deviation for angular hand estimates in the passive localization as a function of those for active localization across all participants (dots). The solid line is the linear line of best fit using an orthogonal distance (or PCA-based) regression which expresses the relationship between the variables without assuming directionality. The dotted line is the unit slope.

### Is variance in movement and hand-estimates related?

Not surprisingly, the precision with which people reached the target with visual feedback correlated with the precision when reaching without feedback (Fig 6A), with a significant shared variance of 22%. The smaller correlation and thus shared variance (6-8%) indicated a weaker relationship between cursor-reach SD and both active and passive hand localization SD (Fig 6B-C). The relationship between no-cursor variance and hand localization variance was even weaker (Fig 6D-E; 3-4%). This suggests there are not many shared processes when estimating the location of the unseen hand and when moving the seen or unseen hand to a target or that additional or different processes dominate in one or the other.

**Figure 6.**
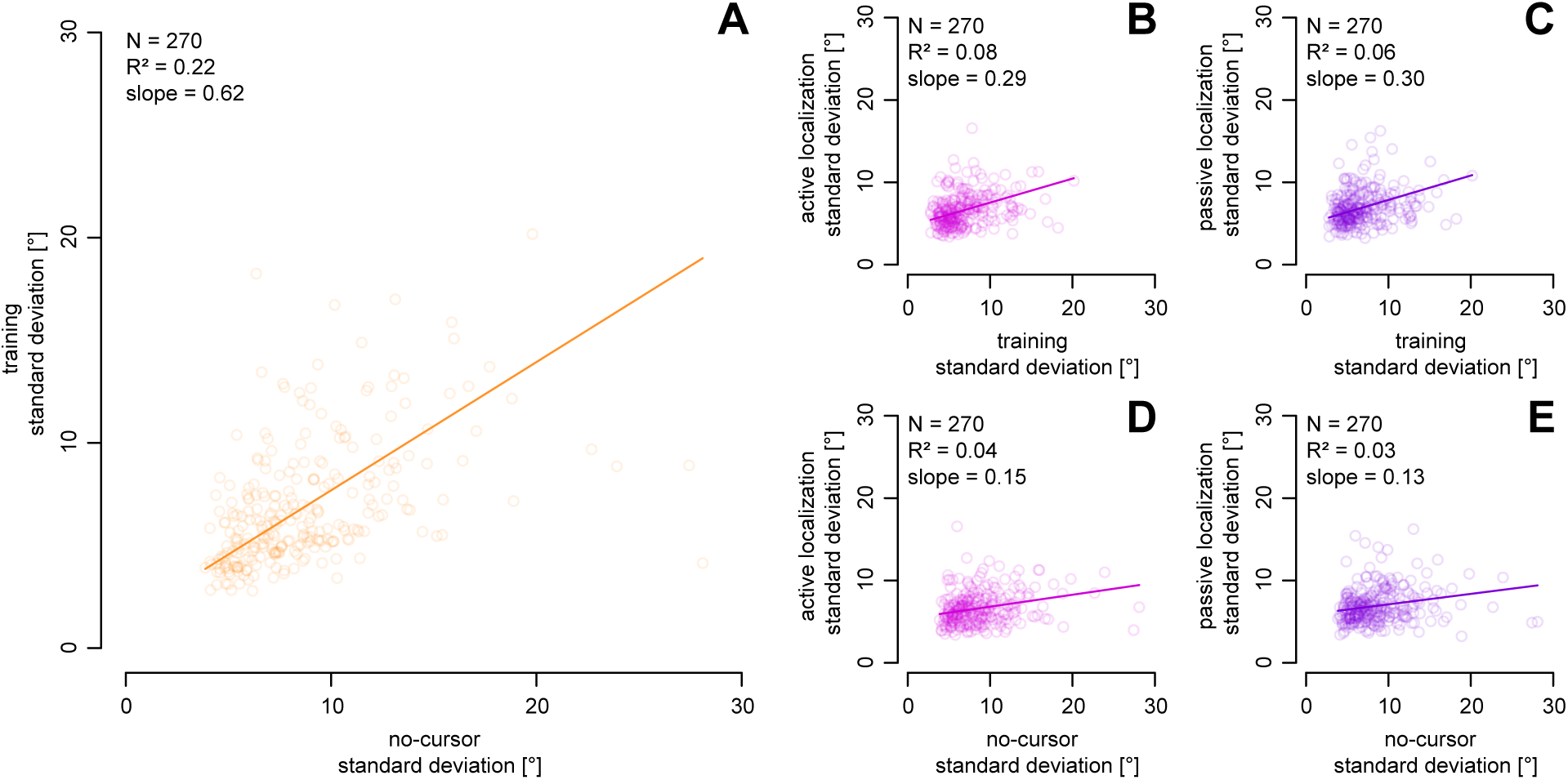
Linear relationship (solid lines) between standard deviation of angular errors for: the aligned-cursor training trials and those for no-cursor trials (**A**) Active localization trials related to aligned-cursor trials (**B**) and no-cursor trials (**D**) and passive localization trials related to aligned-cursor trials (**C**) and no-cursor trials (**E**). Solid lines show the best-fit relationships obtained using orthogonal distance regression (ODR); because ODR minimizes error in both dimensions, the fitted relationship is identical regardless of which variable is placed on either axis. Each dot represents a participant.

### Does motor and sensory variance affect motor adaptation?

As has been previously suggested, greater variance in reaches could facilitate reach adaptation, known as the exploration-exploitation hypothesis, since greater variability may be due to more exploration of the environment that could be exploited for learning. To test this, we measure whether variability in movements prior to cursor-perturbation could also facilitate learning, such as rate of learning, and learning extent (Fig 7), as well as aftereffects (Fig 8) in over 100 participants. We also tested whether this individual motor-variability could predict the magnitude of changes in estimate of unseen hand location that also consistently follows adaptation, or at least after experiencing a visual-proprioceptive mismatch. Reach variance, whether with or without a cursor during the aligned session, did not predict learning rate (Fig. 7A–B), overall learning (Fig. 7E–F), or implicit reach aftereffects (Fig. 8, left column). In short, pre-perturbation reach variability was unrelated to learning or aftereffects.

**Figure 7.**
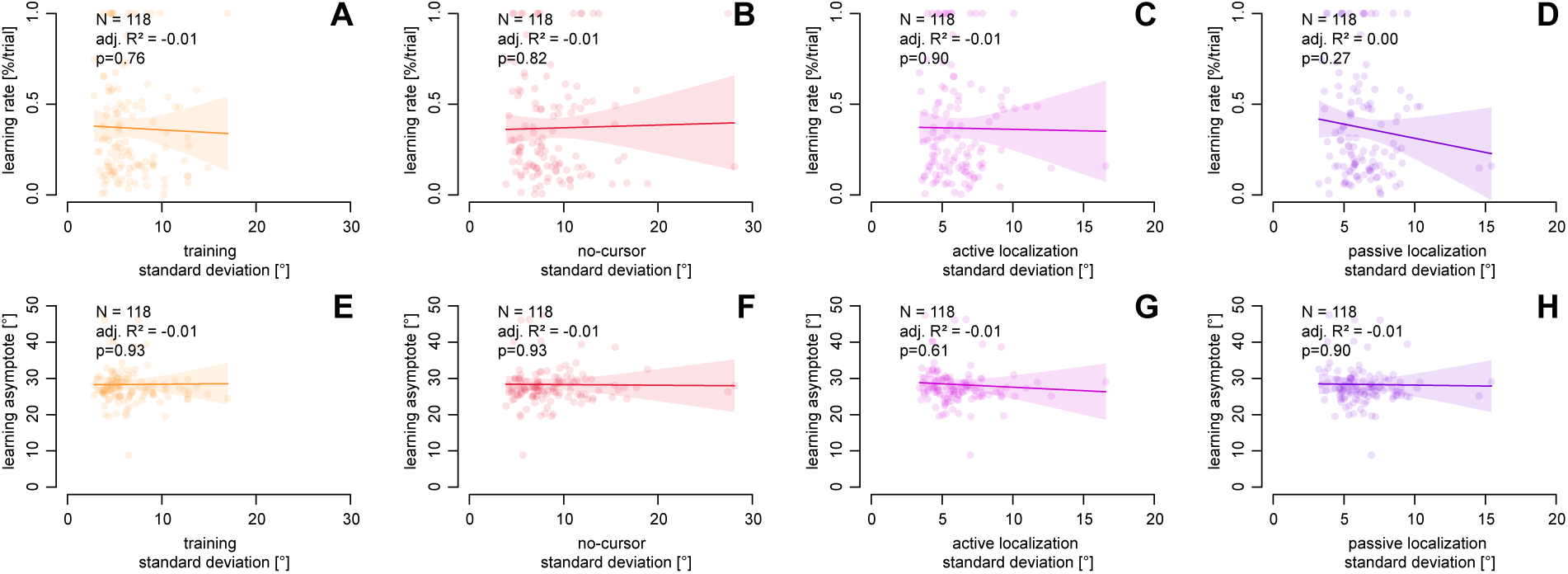
Measures of learning rate (top row) and learning extent (bottom row) as a function of variance for aligned-cursor reaches (**A**, **E**), no-cursor reaches (**B**, **F**), estimates of unseen hand position after active and passive displacement (**C**, **D**, **G**, **H**). Solid lines are lines of best fit. Dots represent each participant.

**Figure 8.**
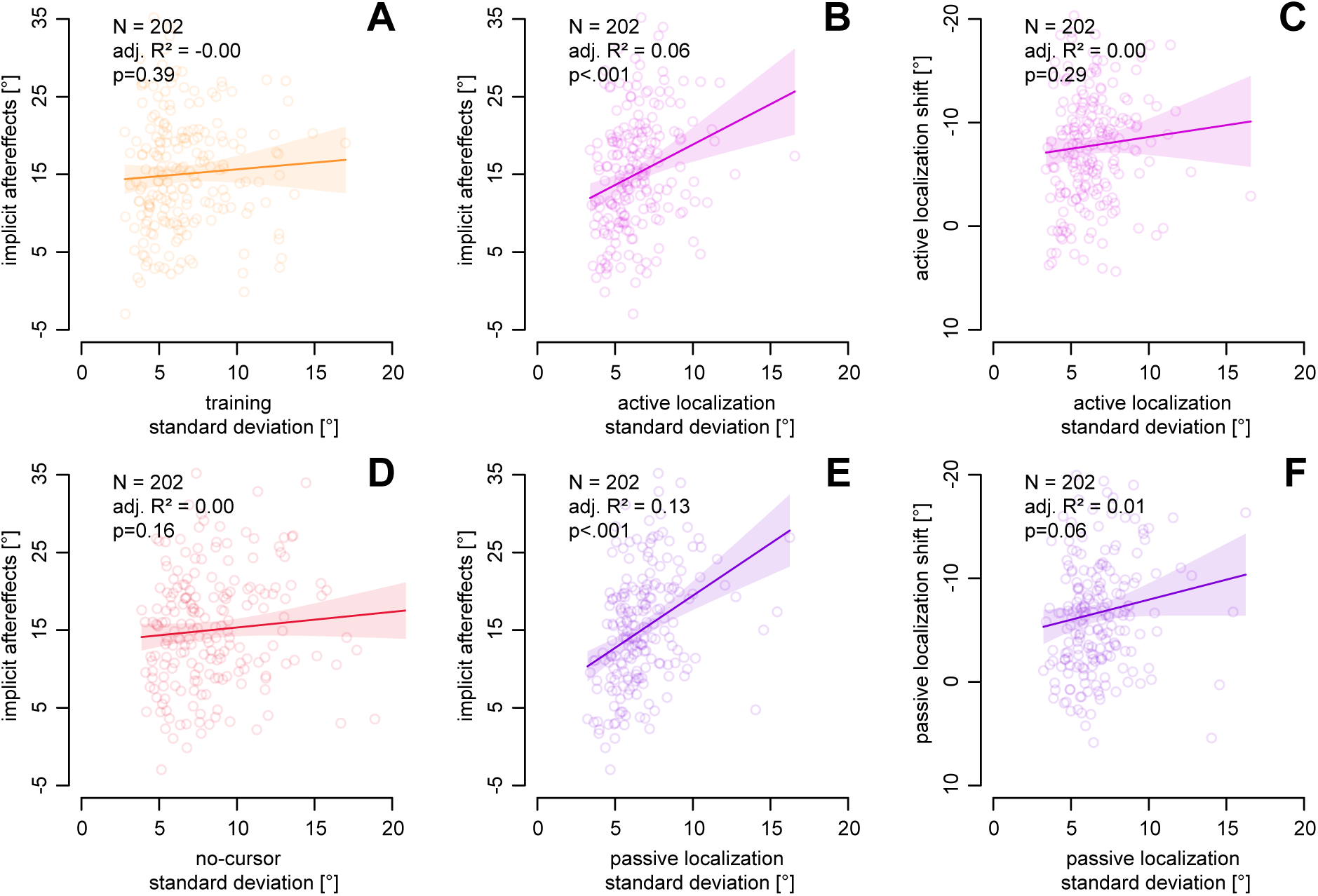
Reach aftereffects as a function of variance in training reaches (**A**) and no-cursor reaches (**D**) and passive and active localization (**B**, **E**). Shift in estimate of unseen hand location as a function of variance in active localization (**C**) and passive localization (**F**). Solid lines are lines of best fit from OLS regression. Each dot represents a participant.

Variance in active and passive localization did not predict learning rate or extent (Fig. 7C–D, G– H), but unexpectedly, it did predict reach aftereffects (Fig. 8, middle column). However, there was no relationship between localization precision and the magnitude of the shift in hand estimates after adaptation (Fig. 8, left column). When considering Fig. 8 as a whole, the only measure of variance that marginally predicted reach aftereffects was hand localization variance. In sum, greater precision in estimating unseen hand position was associated with smaller reach aftereffects, but not with proprioceptive recalibration or motor learning.

In sum, we found that precision in reach movement direction while moving a cursor aligned with the hand did not subsequently predict any feature of learning. However, while variance in hand location estimates can not predict the size of shifts in hand location estimates, it can predict implicit reach aftereffects.

### Relationship between hand localization and reach aftereffects

Not only did the **precision** in hand perception correlate with size of aftereffects, we also found that the magnitude of reach aftereffects was significantly correlated with the magnitude **changes** in hand estimates both for passive and active displacement; with shared variance of 23.6 and 19.6% (Fig 9A,B). Since 30° and 60° training resulted in nearly identical hand-localization shifts (≈5°) and reach aftereffects (≈15°) [37], we treated these conditions as equivalent and included data from both.

**Figure 9.**
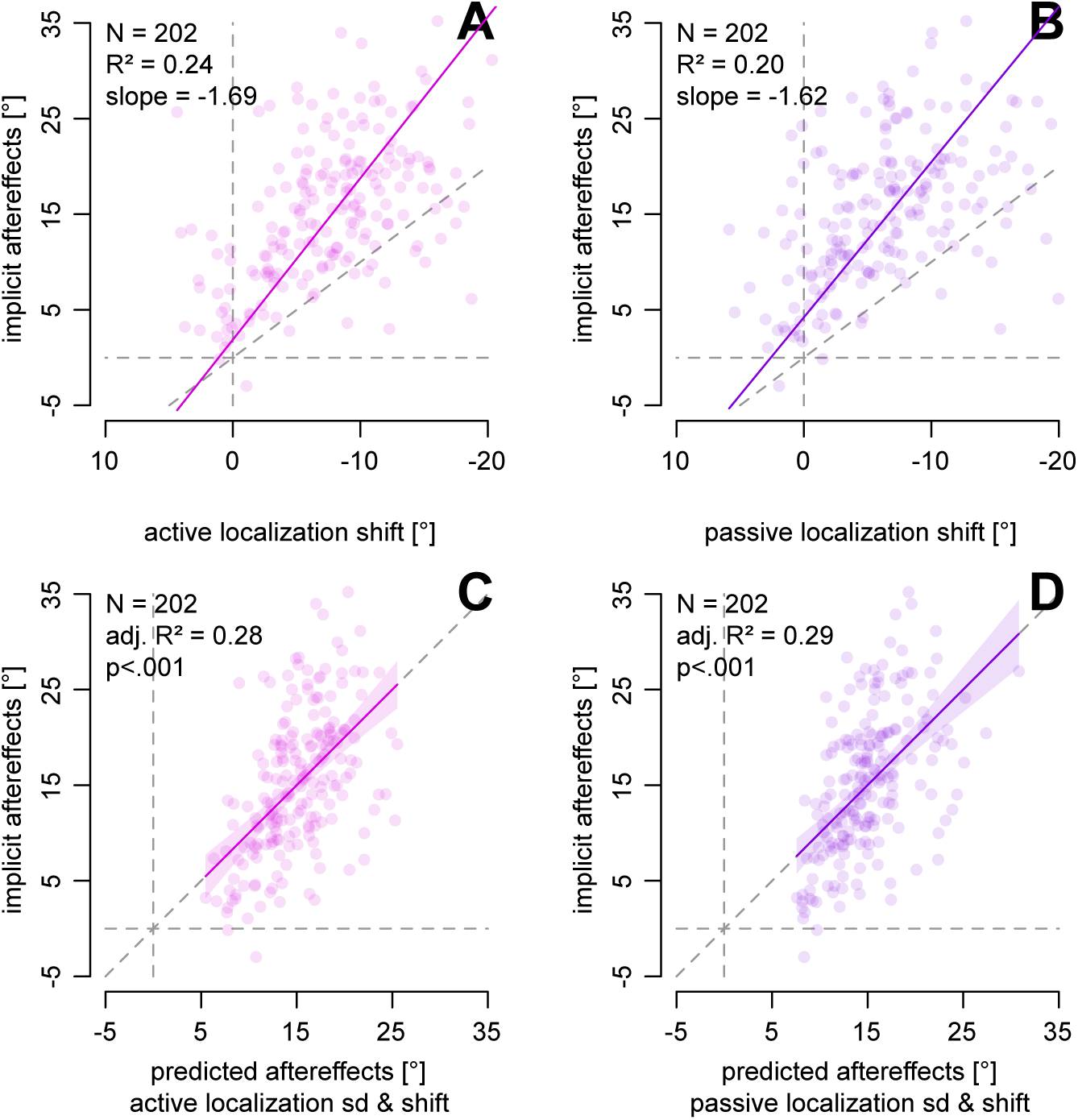
Relationship between magnitude of aftereffects and hand localization measures. **A**,**B**: Localization shifts are related to implicit reach aftereffects. This appears to be equally strong for active (**A**) and passive (**B**) hand localization shifts. **C**,**D**: Implicit aftereffects can be predicted from a combination of hand localization variance in the aligned session and hand localization shifts in the rotated session, for both active (**C**) and passive (**D**) localization.

The variance in aligned hand localization as well as the shift in hand localization after training with rotated feedback are not related to each other (Fig 8C,F) yet each is related with implicit aftereffects (Fig 8B,E; Fig 9A,B). We tested if using both variance and shift in either active or passive hand localization would be better than using only one predictor by comparing their AICs. The AICs of the combined model is better than using just one predictor for both active (AIC: 1332.2 < 1344.2) and passive (AIC: 1330.8 < 1354.6) hand localization (p<0.001). The predicted and observed implicit aftereffects are shown in Fig. 9C–D, and the data fall closely along the unit slope, with R^2^ values of 0.28 and 0.29. This suggests that implicit aftereffects can be reasonably predicted by combining the two hand localization measures (shift and baseline variance). The fact that each variable contributes independently to the prediction may indicate that they reflect distinct subcomponents of implicit motor adaptation. However, this is an exploratory result, so this needs replication. At present we also don’t know what kind of sub-components of implicit reach aftereffects they could each be related to. Primarily, as we and others have found, learning-induced shifts in hand proprioception correlate with aftereffects. Given our previous results, this suggests that part of these implicit motor changes may reflect proprioceptive change in hand localization. How baseline proprioceptive reliability contributes to aftereffects remains an open question.

## Discussion

This study is based on a series of experiments conducted in our lab using the same setup, and similar tasks, where we measured, in separate trials, reaches to visual targets and estimates of the perceived final position of the unseen hand, after the hand movement was self-generated or generated by a robot manipulandum. In each of our studies, we collected a large number of trials to get a reliable estimate of movement and state-estimate *accuracy*. For the current study, we used this large dataset of 270 participants to quantify the *precision* of these movements and hand-estimates. Our within-subjects measures of estimates of the unseen hand location after self-generated movements or robot-generated movements, allowed us to begin exploring the separate contributions of proprioceptive signals and efferent signals on localization. Since the majority of participants then also adapted to a visuomotor rotation, we tested if variance in any of the four measures accounted for individual differences in motor adaptation performance and implicit aftereffects. In summary, this study reveals the remarkable precision of individuals’ estimates of unseen hand position, with minimal influence from additional efferent signals. Moreover, findings indicate that older adults maintain precision in both movements and hand-location estimates compared to younger adults. While motor and sensory noise generally do not predict subsequent visuomotor learning measures, variance and shifts in hand localization unexpectedly predicts the size of reach aftereffects, underscoring the complexity of the relationship between sensory feedback and motor adaptation.

### Proprioceptive and predictive precision

Despite many studies suggesting that proprioceptive estimates of hand position are noisy, our findings suggest that people are relatively accurate and precise when indicating the direction of their unseen dominant hand along the radial axis. This aligns with previous work from our lab and others [7,14,15,45–48], which used a robot manipulandum to position the target hand without visual feedback or a visual goal. Of course, humans are generally less precise when localizing proprioceptive versus visual targets. In our own studies [47,48], reaches to visual targets were 2–3 times more precise than those to proprioceptive hand targets, based on 95% elliptical fits to 2D endpoints. Similarly, Block and Sexton [49] and Liu, Sexton and Block [50] reported comparable levels of endpoint variability when reaching to both visual and proprioceptive targets with an unseen hand. While our current study focused only on directional variance, we observed that no-cursor reaches to a visible target were significantly more variable than estimates of the unseen hand position using the visible left hand. This suggests that although left-hand position is widely used to estimate unseen hand location, the absence of visual feedback during the movement may introduce additional motor noise, potentially confounding localization measures. Therefore, caution is warranted when interpreting such responses as direct reflections of proprioceptive estimates, since they may also include noise introduced by the movement of the indicating limb.

The mixed results regarding how proprioceptive noise varies with age or when compared with visual stimuli may be influenced by other mitigating factors. One such factor is measuring variance or precision necessitates a larger amount of data for reliable results. Many studies reporting variance measures often rely on only a dozen or so data points, which is likely insufficient for a reliable measure. While not critical, certain methods of “placing the hand” may be time-consuming or require extensive exploration. This could introduce additional sources of noise that wouldn’t typically occur when the hand-target is mechanically placed or constrained. Similarly, matching the position of one unseen hand by mirroring it with the other—also unseen —would likely introduce even greater variability than the localization method used in the current study. In some cases, using a larger workspace, outside of where we typically make precise hand movements, could be a factor. Thus, some of the measured variance normally attributed to proprioception may be due to other sources of errors.

The presence of visual cues is one such confounding source of information. Some studies measuring proprioceptive localization also have participants move their target-hand to a visual target prior to having them estimate the position of the unseen hand. However, such visually-guided movements, or even having this second source of information as a cue, would likely dominate the response, reducing the contribution of proprioceptive signals. As a result, such tasks cannot reasonably be considered a measure of proprioception. Thus, in the field’s attempt to rigorously quantify and model proprioceptive acuity, we should be more careful to remove other confounding information sources.

Our series of studies marks a pioneering effort to isolate the predictive contributions to state estimates of end effectors. We achieved this by having participants indicate the location of their unseen hand after directing it to any desired location (active localization), and again in a later block after top-up training where the robot moved their hand to the same locations (passive localization). By comparing the precision of hand localization, we showed that the contribution of efferent signals, beyond what is provided by proprioception, appears to be minimal. Note the target-hand is returned, before localization, so even remembered proprioceptive signals contribute more than efferent–based signals. This holds true not only during baseline conditions but also during visuomotor adaptation, where the resulting recalibration of hand location estimates after active displacement is only slightly larger than for passive. This result is found not only for the studies used in the present dataset but earlier studies in our lab as well [4,9,41], including when the hand-target remains at the target site [4,9]. Since all these studies measured hand localization only after the target hand had stopped moving, the minimal contribution of predictive signals may suggest that such signals are more useful for dynamically monitoring hand position during movement, rather than once the hand has landed. In particular, predictive signals may be more informative—and proprioceptive signals potentially noisier—during the reach itself.

### The effect of age

Another notable finding from this well-powered study is that healthy older adults were no more variable than younger adults when reaching to a visual target—whether or not visual feedback was available—or when localizing their unseen hand. Similarly, recent studies with large samples of older adults have reported little to no difference in reaching or proprioceptive variability, including in wrist position-sense tasks and arm position-matching tasks, when comparing healthy older and younger adults [31,51–55]. Studies that have found that aging leads to poorer precision tend to have small sample sizes (including our own study,[25,26]).

Other methodological drawbacks, as described above, and publication bias may also contribute to purported age-related disparities. The choice of task for estimating hand position can significantly impact the older age group. In one of our prior studies [25,26], we tested a small sample of older adults (N = 9) who estimated the position of their unseen hand relative to a visual landmark using a two-alternative forced-choice (2AFC) task. They found this task somewhat more difficult to understand compared to the older adults in the current dataset (N = 38), who were instead instructed to move their visible indicator hand to the location of their unseen hand. This difference in task comprehension may explain why the uncertainty range (or JND) varied between the two age groups in Cressman et al. [25,26], but not in the current study. It’s worth noting that results using these two methods in young adults do not show any significant differences [47,56]. Being able to see the indicating-hand could mitigate potential increases in noise and age-related differences that occur when individuals cannot see their reaching hand, for both visual and proprioceptive targets (e.g.,[14,15]). Results from the current study support this inference, given that variance in no-cursor reaches to visual targets was 30% larger than variance in reaches made by a visible hand to the unseen target-hand. Seeing the “indicator”—whether a cursor or the opposite hand—may be necessary to avoid introducing additional noise into the task and to obtain a clearer measure of proprioceptive acuity of the “target” hand given the generally higher precision of visual over proprioceptive localization.

Age related differences do exist. Older adults are usually slower at these tasks. Older adults are also poorer than younger adults at sensory tasks that are more cognitively demanding, such as recognizing or reproducing haptic shape [31,51]. Our results suggest that for uncomplicated and unspeeded tasks, motor and proprioceptive acuity doesn’t deteriorate with healthy aging.

### Relation between sensory and motor noise

Our study also allows us to quantify the relationship between noise in visually-guided reaches and hand localization. We observe a clear linear relationship between the variability in closed and open-loop reaches to a visual target despite the greater noise for these open-loop (no-cursor) reaches. We also observe a very strong correlation between the variance measures of passive and active hand localization. This indicates that most of the variance produced when estimating the position of our spontaneously/actively moved hand likely reflects proprioceptive variance, which is captured in the passive hand localization task.

Surprisingly, sensory and motor variability were only weakly correlated with each other, as depicted in Fig 6. Very few other studies have investigated the relationship between proprioceptive and motor variability of the hand, although Saenen et al., [7] found that performance in complicated somatosensory tasks like feeling and reproducing a shape was unrelated to separate measures of motor or proprioceptive function. Overall, our results suggest only a small portion of movement variability reflect somatosensory noise.

### The effect of sensory and motor noise on motor learning

Several theories suggest that variance plays a role in motor adaptation. For instance, the exploration-exploitation hypothesis suggests that motor variance serves a dual role: facilitating exploration to discover new motor strategies and exploiting known strategies to achieve reliable motor performance and adaptation in varying environmental conditions. Likewise, noisier non-visual signals may affect adaptation as greater weight may be placed on the altered visual feedback of the hand. In either case, motor variability or proprioceptive variability should predict aspects of visuomotor learning. However, despite having data from >100 participants, neither motor nor proprioceptive variability could predict the rate or magnitude of learning, as shown in Figure 7. Only variance in hand localization demonstrated any predictive value, specifically for reach aftereffects but only 13% of the variance explained. Motor variability failed to predict aftereffects, and neither type of variance predicted the shift in estimate of hand position that has been shown to consistently emerge with reach adaptation. These results suggest that motor variance does not contribute to visuomotor learning.

However, previous studies from our lab, and others, do show some evidence that the aftereffects and shift in hand-position estimates that occur during visuomotor adaptation or merely experience with a visual-proprioceptive mismatch are related to each other. For a more reliable measure of this possible relationship, we use the current dataset to verify this. Not only did the precision in hand perception correlate with size of aftereffects, we also found that magnitude of reach aftereffects were significantly correlated with changes in hand estimates both for passive and active displacement; with shared variance of 23.6% and 19.6% (fig 7). Moreover these two relationships appear to explain different parts of the variance in reach aftereffects. In summary, as we and others have found, learning-induced shifts in hand proprioception correlate with aftereffects. Given our previous results, this suggests that part of these implicit motor changes may partly reflect proprioceptive change in hand localization.

Lastly, our findings about learning may be limited to adaptation in visuomotor rotation tasks. It remains possible that sensory or motor noise could play a different or more substantial role in adaptation to dynamic perturbations such as force fields.

### Explicit Strategy, Proprioceptive Recalibration, and Aging

Although the present study focuses on sensory and motor precision prior to exposure to a visuomotor perturbation, related work using subsets of this dataset [37–39] has examined how explicit strategy use and age influence different components of visuomotor adaptation, including proprioceptive recalibration. These previous studies of ours show that although explicit instructions—and larger perturbations—can modify initial performance by enabling strategic compensation for the rotation, they do not alter the implicit processes measured after training. In particular, neither proprioceptive recalibration nor shifts in predicted sensory consequences differ between instructed and uninstructed learners, or between small and large visuomotor rotations [37]. Likewise, age-related differences in strategic ability do not translate into differences in implicit learning: older adults may deploy explicit strategies somewhat differently early in training, but they show preserved implicit aftereffects and, notably, exhibit larger proprioceptive recalibration than younger adults [39]. Together, this work indicates that the sensory and motor precision measures emphasized here—assessed prior to the perturbation and dominated by implicit processes—are not influenced by explicit strategy use or by age.

Although explicit strategy use can influence early adaptation, most of the studies contributing to this dataset used a 30° rotation without instruction—a condition that we and others have shown does not elicit meaningful explicit strategies. When instruction or larger perturbations were used, their effects on explicit strategy and learning rate have already been reported in the original publications. Because aim reports and interleaved aftereffects were not collected in these experiments and instructional contexts varied, a full explicit–implicit decomposition is not feasible in the present dataset and may not be tractable even in principle ([57]). Moreover, emerging work in our lab indicates that explicit strategies do not follow a simple exponential time course at the individual level—an assumption underlying many modeling approaches to “learning rate”—further complicating direct comparisons between explicit and implicit processes.

### Informing Future Studies

An additional contribution of the present work is that this unusually large and diverse dataset (https://osf.io/dhk3u/) can serve as a useful empirical resource for *future studies*. Because it provides stable estimates of motor and proprioceptive variability across both younger and older participants, these data can help inform sample-size and trial-count decisions when designing future experiments on adaptation, precision, and sensory recalibration. More broadly, the distributions and confidence intervals reported here provide concrete empirical reference points for variability-based measures, helping design future studies.

## Conclusion

The precision of hand position estimates was as good as that of visually guided reaches. The availability of efferent signals for estimating hand position only reduced variance by 5%, which suggests that estimates of unseen hand position after actively generated reaches largely reflect proprioception, not efference based predictions. Variance in reaches with and without cursor did not predict any measures of adaptation. Implicit reach aftereffects could not be predicted from any reach variance measures before adaptation. However, they could be predicted by the precision of proprioceptive estimates, as well as the magnitude of proprioceptive recalibration after adaptation. This emphasizes the relatively large role that proprioception may play in implicit adaptation.

## References

1. Dijkerman, H. C. & de Haan, E. H. F. Somatosensory processes subserving perception and action. Behav Brain Sci 30, 189–201; discussion 201-239 (2007).

2. Berniker, M. & Kording, K. Estimating the sources of motor errors for adaptation and generalization. Nature Neuroscience 11, 1454–1461 (2008).

3. Bastian, A. J. Learning to predict the future: the cerebellum adapts feedforward movement control. Current Opinion in Neurobiology 16, 645–649 (2006).

4. ’t Hart, B. M. & Henriques, D. Y. P. Separating Predicted and Perceived Sensory Consequences of Motor Learning. PLoS One 11, e0163556 (2016).

5. Vandevoorde, K. & Orban de Xivry, J.-J. Internal model recalibration does not deteriorate with age while motor adaptation does. Neurobiology of Aging 80, 138–153 (2019).

6. Maes, C., Gooijers, J., Orban De Xivry, J.-J., Swinnen, S. P. & Boisgontier, M. P. Two hands, one brain, and aging. Neuroscience & Biobehavioral Reviews 75, 234–256 (2017).

7. Saenen, L., Verheyden, G. & Orban De Xivry, J.-J. The differential effect of age on upper limb sensory processing, proprioception, and motor function. Journal of Neurophysiology 130, 1183–1193 (2023).

8. Ruttle, J. E., ’t Hart, B. M. & Henriques, D. Y. P. Implicit motor learning within three trials. Sci Rep 11, 1627 (2021).

9. Mostafa, A. A., ’t Hart, B. M. & Henriques, D. Y. P. Motor learning without moving: Proprioceptive and predictive hand localization after passive visuoproprioceptive discrepancy training. PLoS One 14, e0221861 (2019).

10. Ghahramani, Z. & Wolpert, D. M. Modular decomposition in visuomotor learning. Nature 386, 392–395 (1997).

11. Ernst, M. O. & Banks, M. S. Humans integrate visual and haptic information in a statistically optimal fashion. Nature 10.1038/415429a (2002) doi:10.1038/415429a.

12. Van Beers, R. J., Sittig, A. C. & Denier Van Der Gon, J. J. The precision of proprioceptive position sense. Experimental Brain Research 122, 367–377 (1998).

13. van Beers, R. J., Wolpert, D. M. & Haggard, P. When Feeling Is More Important Than Seeing in Sensorimotor Adaptation. Current Biology 12, 834–837 (2002).

14. Liu, Y., Sexton, B. M. & Block, H. J. Spatial bias in estimating the position of visual and proprioceptive targets. Journal of Neurophysiology 119, 1879–1888 (2018).

15. Block, H. J. & Sexton, B. M. Visuo-Proprioceptive Control of the Hand in Older Adults. Multisensory Research (2021).

16. Block, H. J. & Bastian, A. J. Sensory reweighting in targeted reaching: effects of conscious effort, error history, and target salience. J Neurophysiol 103, 206–217 (2010).

17. Block, H. J. & Bastian, A. J. Sensory weighting and realignment: independent compensatory processes. J Neurophysiol 106, 59–70 (2011).

18. Jones, S. A. H. & Henriques, D. Y. P. Memory for proprioceptive and multisensory targets is partially coded relative to gaze. Neuropsychologia 10.1016/j.neuropsychologia.2010.10.001 (2010) doi:10.1016/j.neuropsychologia.2010.10.001.

19. Jones, S. A. H., Byrne, P. A., Fiehler, K. & Henriques, D. Y. P. Reach endpoint errors do not vary with movement path of the proprioceptive target. Journal of Neurophysiology 107, 3316–3324 (2012).

20. Reuschel, J., Drewing, K., Henriques, D. Y. P., Rösler, F. & Fiehler, K. Optimal integration of visual and proprioceptive movement information for the perception of trajectory geometry. Exp Brain Res 201, 853–862 (2010).

21. Mikula, L., Gaveau, V., Pisella, L., Khan, A. Z. & Blohm, G. Learned rather than online relative weighting of visual-proprioceptive sensory cues. J Neurophysiol 119, 1981–1992 (2018).

22. Limanowski, J. & Friston, K. Active inference under visuo-proprioceptive conflict: Simulation and empirical results. Sci Rep 10, 4010 (2020).

23. Debats, N. B., Heuer, H. & Kayser, C. Visuo-proprioceptive integration and recalibration with multiple visual stimuli. Sci Rep 11, 21640 (2021).

24. Goble, D. J., Coxon, J. P., Wenderoth, N., Van Impe, A. & Swinnen, S. P. Proprioceptive sensibility in the elderly: degeneration, functional consequences and plastic-adaptive processes. Neurosci Biobehav Rev 33, 271–278 (2009).

25. Cressman, E. K. et al. Proprioceptive recalibration following implicit visuomotor adaptation is preserved in Parkinson’s disease. Exp Brain Res 239, 1551–1565 (2021).

26. Cressman, E. K., Salomonczyk, D. & Henriques, D. Y. P. Visuomotor adaptation and proprioceptive recalibration in older adults. Experimental Brain Research 205, 533–544 (2010).

27. Henriques, D. Y. P., Filippopulos, F., Straube, A. & Eggert, T. The cerebellum is not necessary for visually driven recalibration of hand proprioception. Neuropsychologia 64, 195–204 (2014).

28. Kitchen, N. M. & Miall, R. C. Adaptation of reach action to a novel force-field is not predicted by acuity of dynamic proprioception in either older or younger adults. Exp Brain Res 239, 557–574 (2021).

29. Kitchen, N. M. & Miall, R. C. Proprioceptive deficits in inactive older adults are not reflected in fast targeted reaching movements. Exp Brain Res 237, 531–545 (2019).

30. Vandevoorde, K. & Orban de Xivry, J.-J. Does proprioceptive acuity influence the extent of implicit sensorimotor adaptation in young and older adults? J Neurophysiol 126, 1326–1344 (2021).

31. Plas, S. V. D. & Orban de Xivry, J.-J. Age-related changes in proprioception are of limited size, outcome-dependent and task-dependent. bioRxiv 2025.05.22.655043 (2025) doi:10.1101/2025.05.22.655043.

32. Dhawale, A. K., Smith, M. A. & Ölveczky, B. P. The Role of Variability in Motor Learning. Annu. Rev. Neurosci. 40, 479–498 (2017).

33. Van Beers, R. J. Motor Learning Is Optimally Tuned to the Properties of Motor Noise. Neuron 63, 406–417 (2009).

34. Wei, K. & Körding, K. Uncertainty of feedback and state estimation determines the speed of motor adaptation. Front Comput Neurosci 4, 11 (2010).

35. Tsay, J. S., Kim, H. E., Parvin, D. E., Stover, A. R. & Ivry, R. B. Individual differences in proprioception predict the extent of implicit sensorimotor adaptation. Journal of Neurophysiology 125, 1307–1321 (2021).

36. He, K. et al. The Statistical Determinants of the Speed of Motor Learning. PLoS Comput Biol 12, e1005023 (2016).

37. Modchalingam, S., Vachon, C. M., ’t Hart, B. M. & Henriques, D. Y. P. The effects of awareness of the perturbation during motor adaptation on hand localization. PLoS One 14, e0220884 (2019).

38. Gastrock, R. Q., Modchalingam, S., ’t Hart, B. M. & Henriques, D. Y. P. External error attribution dampens efferent-based predictions but not proprioceptive changes in hand localization. Sci Rep 10, 19918 (2020).

39. Vachon, C. M., Modchalingam, S., ’t Hart, B. M. & Henriques, D. Y. P. The effect of age on visuomotor learning processes. PloS One 15, e0239032 (2020).

40. Clayton, H. A., ’t Hart, B. M. & Henriques, D. Y. P. Sensing hand position in Ehlers-Danlos syndrome. Somatosensory & Motor Research 38, 303–314 (2021).

41. Ruttle, J. E., ’t Hart, B. M. & Henriques, D. Y. P. Implicit motor learning within three trials. Sci Rep 11, 1627 (2021).

42. Rouder, J. N., Morey, R. D., Speckman, P. L. & Province, J. M. Default Bayes factors for ANOVA designs. Journal of Mathematical Psychology 56, 356–374 (2012).

43. Rouder, J. N., Speckman, P. L., Sun, D., Morey, R. D. & Iverson, G. Bayesian t tests for accepting and rejecting the null hypothesis. Psychon Bull Rev 16, 225–237 (2009).

44. D’Amario, S., Ruttle, J. E., Hart, B. M. ’t & Henriques, D. Y. P. Implicit Adaptation is Fast, Robust and Independent from Explicit Adaptation. 2024.04.10.588930 Preprint at 10.1101/2024.04.10.588930 (2024).

45. Jones, S. A. H. & Henriques, D. Y. P. Memory for proprioceptive and multisensory targets is partially coded relative to gaze. Neuropsychologia 10.1016/j.neuropsychologia.2010.10.001 (2010) doi:10.1016/j.neuropsychologia.2010.10.001.

46. Jones, S. A. H., Byrne, P. A., Fiehler, K. & Henriques, D. Y. P. Reach endpoint errors do not vary with movement path of the proprioceptive target. Journal of Neurophysiology 107, 3316–3324 (2012).

47. Jones, S. A. H., Fiehler, K. & Henriques, D. Y. P. A task-dependent effect of memory and hand-target on proprioceptive localization. Neuropsychologia 50, 1462–1470 (2012).

48. Jones, S. A. H., Cressman, E. K. & Henriques, D. Y. P. Proprioceptive localization of the left and right hands. Experimental Brain Research 204, 373–383 (2010).

49. Block, H. J. & Sexton, B. M. Visuo-Proprioceptive Control of the Hand in Older Adults. Multisensory Research (2021).

50. Liu, Y., Sexton, B. M. & Block, H. J. Spatial bias in estimating the position of visual and proprioceptive targets. Journal of Neurophysiology 119, 1879–1888 (2018).

51. Saenen, L., Verheyden, G. & Orban De Xivry, J.-J. The differential effect of age on upper limb sensory processing, proprioception, and motor function. Journal of Neurophysiology 130, 1183–1193 (2023).

52. Vandevoorde, K. & Orban de Xivry, J.-J. Internal model recalibration does not deteriorate with age while motor adaptation does. Neurobiology of Aging 80, 138–153 (2019).

53. Vandevoorde, K. & Orban de Xivry, J.-J. Does proprioceptive acuity influence the extent of implicit sensorimotor adaptation in young and older adults? J Neurophysiol 126, 1326–1344 (2021).

54. Maes, C., Gooijers, J., Orban De Xivry, J.-J., Swinnen, S. P. & Boisgontier, M. P. Two hands, one brain, and aging. Neuroscience & Biobehavioral Reviews 75, 234–256 (2017).

55. Parthasharathy, M., Mantini, D. & Orban De Xivry, J.-J. Increased upper-limb sensory attenuation with age. Journal of Neurophysiology 127, 474–492 (2022).

56. Jones, S. A. H., Cressman, E. K. & Henriques, D. Y. P. Proprioceptive localization of the left and right hands. Experimental Brain Research 204, 373–383 (2010).

57. ’t Hart, B. M., et al. Measures of Implicit and Explicit Adaptation Do Not Linearly Add. eNeuro 11, ENEURO.0021-23.2024 (2024).

